# The ecological dynamics and consequences of phytoplasma infection in the spring ephemeral white trillium

**DOI:** 10.1101/2025.10.06.680565

**Authors:** Julia A. Boyle, Yanggaolan Yang, Amalia Caballero, Eleanor N. Hector, Alex Ellett, Emily Glasgow, John R. Stinchcombe, Megan Bontrager

## Abstract

Ecologists have long sought to link within-population dynamics to patterns of species occurrence and persistence across the range. Theoretical models suggest pathogens have important implications for plant populations and perhaps even limit plant distributions, though more empirical evidence is needed, especially in wild plant populations. In the field, we characterized the infection dynamics of phytoplasma, a vectored and sterilizing bacterial pathogen, in wild populations of white trillium. We identified the highest molecular matches for the local bacterial strain and putative vectors. We then leveraged iNaturalist data to reveal patterns and potential drivers of phytoplasma occurrence across the range of white trillium. Within populations, we found that phytoplasma symptoms were more likely in low host density plots, where insects occupied a higher proportion of trillium, indicative of an encounter-dilution effect. Leafhoppers in the genus *Empoasca* were positive for phytoplasma, suggesting they are a candidate vector of disease between trilliums at our field site. The closest match to our local phytoplasma was related to ‘*Candidatus* Phytoplasma pruni’, ribosomal group 16SrIII-F. Phytoplasma symptoms were widely distributed across trillium’s range, even at range edges, and probability of symptoms was significantly increased by proximity to cropland. We studied the impacts of a sterilizing, vector-transmitted pathogen on host plant populations and patterns of infection across host range. Our results suggest sap-feeding insects do not occupy all hosts at high host densities, thereby reducing transmission and leading to higher likelihood of finding infection at low host densities, and a higher proportion of host plants infected at low host densities. At a range-wide scale, we found evidence that the pathogen was able to persist across both the core and edge of host ranges. Thus, negative-density dependent sterilizing pathogens may challenge host populations across the range.

## Introduction

Understanding how within-population dynamics scale up to affect species’ metapopulation dynamics and geographic distributions has been a central goal of ecological research for decades (Harrison and Taylor, 1997; Freckleton et al., 2005; Oldfather and Ackerly, 2019) and understanding the impacts of pathogens on these large-scale processes remains challenging (Gilbert 2002). Plant populations display density-dependent dynamics in interactions with competitors, pollinators, seed predators, herbivores, and others (Gilbert 2002; Gunton and Kunin, 2009; Kim et al., 2013; Hegland 2014). The addition of phytopathogens may reinforce or disrupt a plant population’s density-dependent processes (Parker and Gilbert, 2018), which could lead to altered distribution patterns or demographic structure. Much of our understanding of plant pathogens comes from agricultural systems, but natural populations may have fundamentally different infection dynamics due to environmental and demographic heterogeneity (Alexander 2010). Disease dynamics in natural populations are also important because populations can act as reservoirs of pathogens (Webster et al., 2007; Petersen et al., 2019) and because disease outbreaks are a conservation concern, even for abundant plant species (Frankel 2008; Pautasso et al., 2013).

Pathogen success hinges on the properties of host populations, and host population density often plays an important role at a local scale (Burdon and Chilvers, 1982). Density-dependent dynamics occur when transmission has a linear relationship with the absolute number of infected individuals (Antonovics 2017); positive density-dependence has been observed in many wild plant populations, often when pathogens are transmitted aerially or via direct contact (Carlsson and Elmqvist, 1992; Gilbert et al., 1994; Lively et al., 1995; Bell et al., 2006). Density-dependent transmission may result in different proportions of individuals infected at different host densities. In contrast, frequency-dependent dynamics occur when transmission increases with the proportion of hosts that are infected, instead of correlating with host density (Antonovics 2017; Antonovics et al., 2023). Over time, this may result in similar proportions of individuals infected across populations of differing density. While vector-transmitted pathogens are often considered frequency-dependent, they may possess qualities of both density-dependent and frequency-dependent transmission because transmission can initially increase as host density increases, but decrease at higher host density (Antonovics et al., 1995; 2017; 2023). Transmission may decrease at high host density because vectors are limited by time, satiation, or abundance, reducing the chance that a host interacts with a vector; this phenomenon is known as the encounter-dilution effect (Mooring and Hart, 1992; Krebs et al., 2014). Recently, Antonovics and colleagues (2023) found that pollinators were more effective transmission vectors of anther-smut fungus at lower host densities compared to higher host densities, providing evidence of the encounter-dilution effect in plant-vector-pathogen interactions at relatively small spatial scales.

Scaling up phytopathogen dynamics from a local host population to explore patterns across a whole host range is empirically challenging and therefore, rarely done (Bruns et al., 2019 *a*; Uricchio et al., 2023). Species ranges are often constrained by the climatic and resource parameters a species can tolerate (Lee-Yaw et al., 2016), and it remains an open question whether species interactions frequently expand or limit ranges and abiotic niches (Afkhami et al., 2014; Harrison et al., 2018; Uricchio et al., 2023; Nathan et al., 2023). Classic models with host-specific pathogens predict that regions of low host density, such as some range edges, should also have low pathogen loads due to positive density-dependence, forming a ‘disease-free halo’ (Antonovics 2009; Bruns et al., 2019 *a*). However, frequency-dependent pathogens may be found even in areas of low host density. Bruns et al. (2019 *a*) found that a vectored phytopathogen, anther-smut disease, was maintained at a host’s range limits and occurred at higher prevalence at range edges compared to range centres. Further work indicates that this pathogen constrains the range of a host species and may eventually lead to host range contraction or extinction (Uricchio et al., 2023). The persistence of plant populations at range edges is important for the ability of a species to shift ranges and respond to climate change (Rehm et al., 2015; Hargreaves and Eckert, 2019), thus necessitating a deeper understanding of how pathogens affect plant populations across host ranges.

Almost 150 years ago, Erwin Smith described a “monstrous form” of *Trillium grandiflorum* (Smith 1879). Instead of the characteristic white blooms, Smith observed trillium with virescent petals and more numerous leaves held on longer petioles. While Smith suggested these plants might be taxonomically distinct entities, we now know that these morphological variations reflect symptoms of phytoplasma infection (Hooper et al., 1971). Phytoplasmas are bacterial pathogens that live obligately in the intracellular phloem tissue of plants and the bodies of their insect vectors, which are phloem-feeding insects in the order Hemiptera (Trivellone 2019; Weintraub et al., 2019; Trivellone and Dietrich 2021; Pedroncelli et al., 2026). Insect vectors are exposed to phytoplasmas when they ingest the phloem of infected plants (Christensen et al., 2005). Phytoplasmas have been found in over 1000 plant species including those used as timber, ornamental plants, and crops (Lee et al., 2000; Kumari et al. 2019). As a result of flower abnormalities, causing petals to develop as leaf-like structures, phytoplasmas can sterilize their plant hosts (Lee et al., 2000; MacLean et al. 2014). Infection is systemic and lifelong (Lee et al., 2000; Wright et al., 2022; Ustun et al., 2023). The longevity of infection may be especially consequential for populations of susceptible perennial plants, as it means these individuals may act as a source of infection for many years. *Trillium grandiflorum* is one such long-lived species; individuals become reproductive after 7-10 years, with a total lifespan of over 30 years (Hanzawa and Kalisz, 1993; Lamoureux and Larose, 2002). Theoretical models predict that sterilizing pathogens in small and long-lived host populations can drive host populations to extinction more often than lethal pathogens (Anderson & May, 1981; Boots & Sasaki, 2001; Gerber et al., 2005; Antonovics 2009). Thus, phytoplasma may have grave consequences for the persistence of trillium and other susceptible plants’ populations. Smith’s (1879) report of symptoms resembling a trillium-dwelling phytoplasma was in Michigan, USA, suggesting this species interaction has long been established in northeastern America. Although the first Canadian confirmation of phytoplasma infection in trillium occurred in the last decade (closely matching ‘*Candidatus* Phytoplasma pruni’, ribosomal 16SrIII-F group, Arocha-Rosete et al., 2016), observations of the symptoms of phytoplasma infection in *Trillium* in Ontario were documented long before (Gates 1917). The phenotypic, fitness, and population-level effects of phytoplasma on trillium have never been quantified, thus characterizing these effects is one step towards determining if this pathogen poses a threat to trillium persistence or if they are stably co-existing.

We explored the links between phytoplasma symptoms and the performance, stage structure, and density of wild populations of the spring ephemeral *Trillium grandiflorum*. We further characterized the strain of phytoplasma and possible insect vectors at our field site. We then leveraged iNaturalist data to examine how patterns in phytoplasma symptoms within plant populations scale up across the range of *T. grandiflorum*. We hypothesized that phytoplasma would reduce trillium fitness (MacLean et al. 2014) and reduce the relative abundance of earlier demographic stage classes within populations. Given that phytoplasma is vector transmitted, we hypothesized that its transmission would be primarily frequency-dependent (Antonovics 2017; Antonovics et al., 2023), resulting in an equal chance and rate of infection in populations regardless of host density. However, we had an alternative hypothesis that the encounter-dilution effect could reduce transmission at high host densities due to vector behaviour or limitation (Mooring and Hart, 1992; Krebs et al., 2014; Antonovics et al. 2023), resulting in a lower proportion of infected hosts, and more consistent with negative density-dependence. Finally, we predicted that phytoplasma infection would be prevalent even at range edges due to frequency-dependent, vector-mediated transmission (Bruns et al. 2019 *a*). Our work characterizes infection dynamics of a fearsome and understudied plant pathogen in a wild host population and across the host’s range.

## Methods

### Study organisms and site

*Trillium grandiflorum* (Michx.) Salisb., white trillium, is a spring ephemeral native to the deciduous woodlands of eastern North America. Its range spans from Minnesota to the Eastern coast, and from Georgia to northern Ontario (Case 1993). *Trillium grandiflorum* reproduces by seed and clonal reproduction by rhizomes is rare (Ohara 1989); seeds of this species are frequently dispersed by ants over short distances (Kalisz et al., 1999). Trillium have four distinct demographic stages: a seedling stage, a single leaf vegetative stage, a three-leaf vegetative stage, and a reproductive stage (Ohara 1989; Rooney and Gross, 2003). Once a trillium has first bloomed it often continues to bloom annually but can go dormant for one or more years (persisting as a rhizome with no aboveground structures) or revert to a three-leaf non-reproductive stage. In Ontario, white trillium begins flowering in mid-May and sets seed by July, after which the aboveground parts of the plant die back. There are large populations of wild *T. grandiflorum* at our study site (Figure S2), the Koffler Scientific Reserve (KSR, www.ksr.utoronto.ca) in Ontario, Canada (44°01’48”N, 79°32’01”W). Symptoms of phytoplasma infection have been observed in trillium populations at KSR since at least the early 2000s (A. Weis, personal communication). Phytoplasma infection is most apparent when the white trilliums are flowering, as petals will range from having a streak of green to full virescence or phyllody (Figure 1). The phytoplasma previously identified in a different *T. grandiflorum* population was a 99% sequence identity match to ‘*Candidatus* Phytoplasma pruni’, ribosomal group 16SrIII-F (Arocha-Rosete et al., 2016).

**Figure 1.**
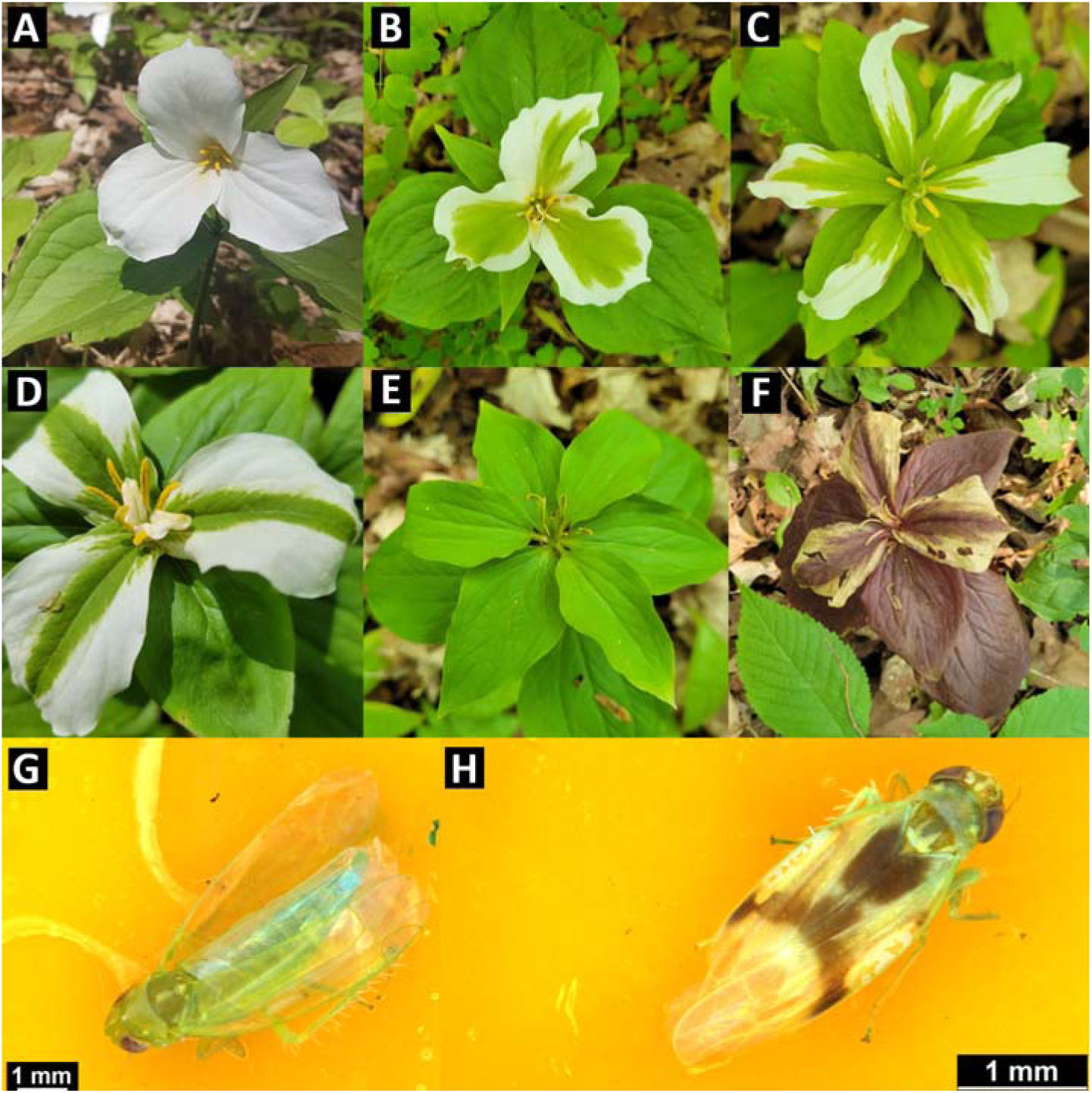
Diversity of phytoplasma infection symptoms in *Trillium grandiflorum* compared to an asymptomatic plant (A). Symptoms include virescence of petals (B-D), larger sepals (C-F), phyllody (E), as well as other deformations including extra petals (C) and the absence of pollen (E, F). Discolouration in an infected plant late in the flowering season (F). Using sticky traps, we identified *Empoasca* spp. leafhoppers positive for phytoplasma (G, H). All photos by authors, images of *T. grandiflorum* were taken at the Koffler Scientific Reserve, ON, Canada. Alt text: A multi-panel figure of photographs. The first six are trillium showing diverse symptoms of phytoplasma infection, while the last two images are of leafhopper morphotypes that were positive for phytoplasma infection.

### Identification of potential insect vectors and phytoplasma strain

To identify the phytoplasma present in trillium, we sampled a leaf from a flowering plant showing virescence symptoms and extracted DNA using QIAGEN’s DNeasy® PowerSoil® Pro Kit. Then, we used sticky traps to broadly sample the phloem-sucking insect community interacting with and near trillium, with the goal of identifying local phytoplasma vectors. We deployed yellow sticky traps (15.24 cm x 25.4 cm) attached to bamboo poles just above trillium height at 0.5 m above ground (Figure S2). We placed 20 traps across the sampled KSR trillium regions on June 6th 2023, collected them after 6 days, and stored traps at -20. We counted and collected potential vectors (e.g. phloem-sucking insects) from each trap, and categorized them into morphotypes. We chose 110 insects across potential vector morphotypes and extracted their DNA using QIAGEN’s DNeasy® Blood & Tissue Kit, following the protocol for insect specimens. We detected and amplified the 16S rRNA region of phytoplasma in insect and plant samples with a nested polymerase chain reaction (PCR) using two universal phytoplasma primer pairs R16mF2/R1 and R16F2n/R2 (primers and PCR conditions as reported in Gundersen & Lee, 1996) and visualised the presence of phytoplasma using gel electrophoresis. We amplified the mitochondrial cytochrome oxidase subunit 1 region of insect samples positive for phytoplasma using the lib220-1/LepR primers (Hajibabaei et al., 2006). The reactions used 1 µL template, 1 µL each of forward and reverse primers, 12.5 µL GoTaq® Master mix (Promega) and 9.5 µL nuclease-free water). Insect cycling conditions were as follows: 94 for 60s, then 5 cycles of 94 for 40s, 45 for 40s, and 72 for 60s, followed by 35 cycles of 94 for 40s, 51 for 40s, 72 for 60s, and a final extension of 72 for 5 mins. We cleaned all PCR products using 0.2 ul Exonuclease I (New England Biolabs, 20U/ul) and 0.2ul calf intestinal alkaline phosphatase (New England Biolabs,10U/ul). Then, to identify the phytoplasma present in plants and the insects that harbour it, we performed Sanger sequencing on PCR products (at the Centre for Analysis of Genome Function and Evolution, University of Toronto) and identified the most similar sequences using the National Centre for Biotechnology Information’s nucleotide basic local alignment search tool using the megablast algorithm (Altschul et al., 1990; Chen et al., 2015).

### Effects of phytoplasma infection on *Trillium grandiflorum* growth and reproduction

To investigate whether infection was associated with differences in plant size, we quantified total plant height, leaf length, and sepal length in symptomatic (n = 32) and asymptomatic (n = 30) trillium during peak flowering season (May 20-29, 2023) across 30 locations in an approximately 15-hectare section of mixed deciduous forest at KSR. We determined infection status by visually inspecting petals for virescence and phyllody; we suspect that some infected individuals have no visible symptoms, so this method likely underestimates the overall infection rate of the population. To test for differences in plant size phenotypes associated with infection, we used linear mixed effects models with infection status as a predictor and location as a random effect. We applied a natural log transformation on responses as needed to improve normality. For statistical analysis, we used R v4.2.0 (R Core Team 2022), with the *tidyverse* (Wickham et al., 2019), *lme4* (Bates et al., 2015), and *lmerTest* (Kuznetsova et al., 2017) packages. All linear models were corrected using a type III ANOVA with the *car* package (Fox and Weisberg Sanford, 2019). All data and code are archived on Dryad (Boyle et al., 2026 *a*).

To quantify differences in seed production due to infection, we first characterized the relationship between size and seed number in asymptomatic trilliums. We haphazardly sampled 100 individuals on 13 June 2024, aiming to maximize variation in plant size (Figure 2A). On these plants we measured leaf length and seed set, then estimated the relationship between them using a generalized linear model with a Poisson family. Then, we counted the number of seeds produced by 81 haphazardly chosen symptomatic trillium in early to mid-June 2024. To estimate potential reproductive loss due to infection in a given environment, we identified the nearest asymptomatic trillium of similar size and estimated its predicted seed count (rounded to the nearest whole number) from the relationship between leaf length and seed number using the predict.glm function; logistical and time constraints prevented us from also sampling seeds from these asymptomatic individuals and determining this relationship directly. Then, we used a negative binomial generalized linear mixed model with seed count as a response, symptom status as a fixed effect, and pair number as a random effect.

**Figure 2.**
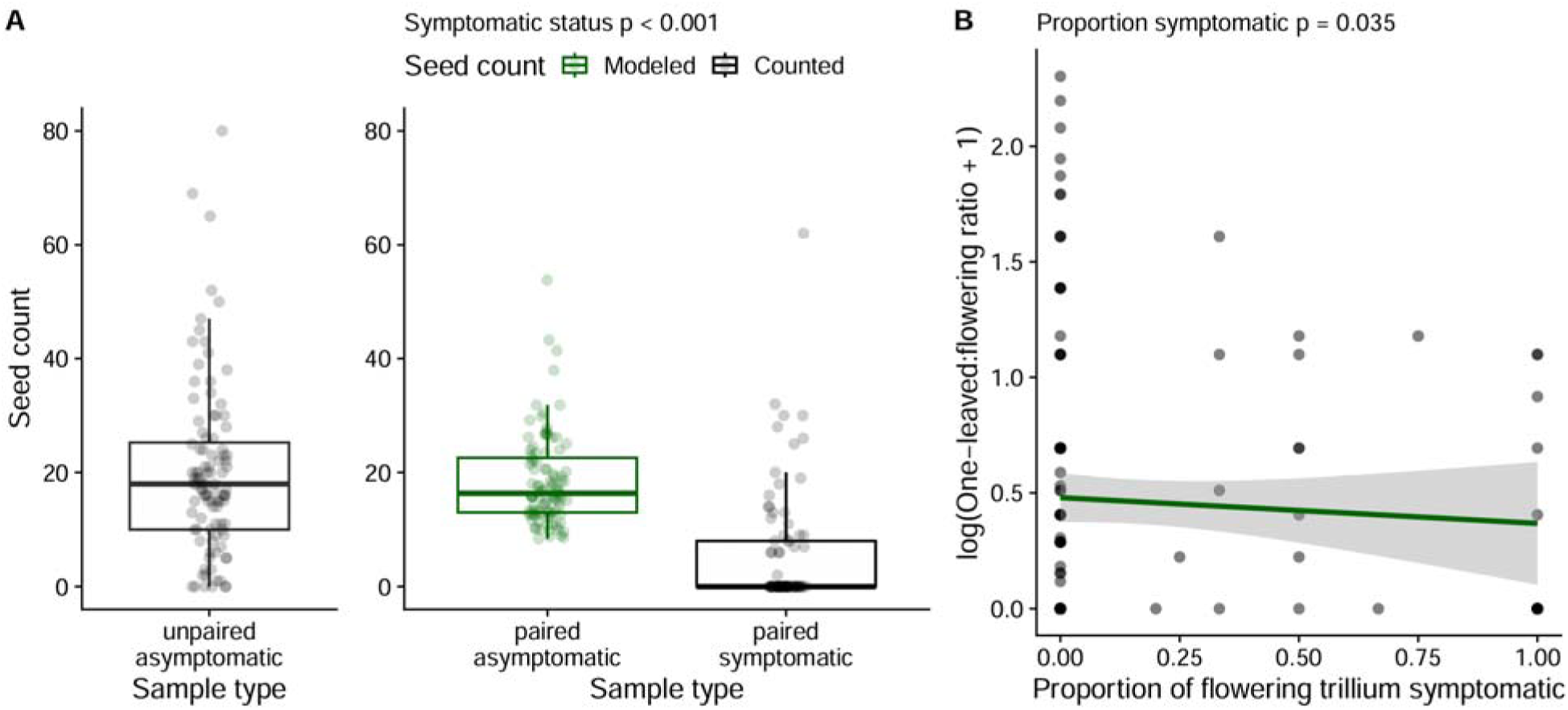
Relationship between phytoplasma symptoms, trillium reproduction, and the relative abundance of the one-leaved stage class. (A) Using the relationship between leaf size and seed set in asymptomatic trillium (black ‘unpaired asymptomatic’ samples), we found that counted symptomatic seed set was significantly less than nearby modelled asymptomatic seed set (*X*^2^ = 34.7, *p* < 0.001). The boxplots show the median with the lower and upper hinges corresponding to the 25th and 75th percentiles, and the upper and lower whiskers represent the largest or smallest value, respectively, that is no further than 1.5 times the inter-quartile range from the hinge. Individual data points represent either direct seed counts (black “counted”) or seed counts predicted based on an empirical relationship between leaf size and seed set (green “modeled”); data points are shown at lower opacity superimposed on the boxplots. (B) In 2024, the ratio of one-leaved trillium to flowering three-leaved trillium declined slightly but significantly across plots with higher infection rates. The scatterplot shows data points and a linear model fit. Alt text: A two-panel figure. The first panel is a boxplot showing reduced seed set in symptomatic trillium, while the second panel is a scatterplot showing a reduced ratio of one-leaved to flowering three-leaved trillium in plots with more infection.

### Within-population infection dynamics

To determine the outcome of phytoplasma transmission at varying host densities within a population, we conducted surveys of trillium density and phytoplasma symptom frequency in the *T. grandiflorum* population of KSR (Figure S3) in mid-late May 2018 and 2023. In both survey years, 1 m^2^ plots were chosen haphazardly by throwing a flag into the trillium population (Figure S2); all plots in a given year were at least 5 m apart. We grouped subsampling locations on geographic features within the larger population (e.g., separate hill slopes). In 2018, we sampled 210 plots across 21 subsample regions; we counted the number of flowering *T. grandiflorum* in the plot and used petal virescence and phyllody as a proxy for their infection status. In 2023, we sampled 64 additional plots across 8 subsample regions with the aim of including more high-density populations and surveying potential vectors. We counted the number of non-flowering and flowering *T. grandiflorum* individuals that had at least three leaves, noted the infection status of flowering trillium, and noted whether phytophagous Hemiptera juveniles were present on the plants. We noted the presence of Hemiptera juveniles for several reasons. Firstly, the presence of Hemiptera juveniles were a direct result of adult visitation and thus was used as a proxy for adult sap-feeding insect attraction, visitation, and feeding on the plant. Phytoplasma is transmitted from vector to host plant by phloem-feeding on the plant (Weintraub et al. 2019). Secondly, Hemipteran juveniles were flightless and so we were able to accurately record their presence, which would have been more challenging with potential adult insect vectors. It is important to note that Hemipteran juvenile presence was not differentiated to species and instar stage due to sampling and expertise constraints, and so it represents a survey of the attraction of both phytoplasma vectors and non-vectors-as such, our results may be underestimating the strength of the relationship between phytoplasma vector presence and infection.

Across both surveys, we estimated the proportion of trillium symptomatic by dividing the number of symptomatic flowering plants by the total number of flowering trilliums (we used only flowering individuals in the denominator because infection was determined by visual petal symptoms). In addition, using data from our 2023 survey, we calculated the proportion of trillium with phytophagous Hemipterans present by dividing the number of trilliums with Hemiptera by the number of trilliums in the plot; we did this for total trilliums and also for only flowering trillium.

To test whether the number of symptomatic trilliums was dependent on the number of flowering trilliums in a patch and sampling year, we used zero-inflated Poisson models with (*GLMMadaptive*; Rizopoulos 2025) and without (*pscl*; Zeileis et al., 2008) a random effect of subsampling location. These zero-inflated models predict both the zeros (infection presence or absence), and the counts (number of symptomatic trillium) separately. We compared the log likelihood of models with and without random effects and used the better fitting model. Then, using data from 2023 only, we used the same models without the year predictor and added the proportion (values between 0-1) of flowering trillium with Hemiptera as an additional predictor. Hemipterans may also be attracted to non-flowering trillium, and the number of flowering trillium was correlated with the number of non-flowering trillium (*cor* = 0.571, *t* = 5.53, *df* = 63, *p* < 0.001 using the cor.test function). To further explore patterns of trillium density and potential vector occurrence in the 2023 data, we used a binomial generalized linear model with the proportion of trillium with Hemiptera as the response and the total trillium density as the predictor and a weight. Additionally, we used a binomial generalized linear model to test whether the proportion of symptomatic flowering trillium in a plot was associated with the proportion of flowering trillium with Hemiptera on them, weighted by total trillium density.

If phytoplasma infection affects seed set then populations may recruit fewer new plants, altering the age class distribution of surrounding plants and challenging long-term population persistence. To determine whether phytoplasma symptoms affected recruitment and the age class distribution of KSR trillium populations, we first calculated plot level ratios of vegetative to reproductive trillium. We used ratios as responses because they account for shifts in underlying flowering trillium density. Using data from the 1 m^2^ plots surveyed in 2023, we calculated the natural log of the ratio of vegetative to flowering three-leaved trillium. Separately, in 2024, we sampled 0.5 m^2^ plots centered around the asymptomatic and symptomatic individuals measured for seed production, and recorded the number of one-leaved, three-leaved vegetative, and three-leaved flowering trillium. We calculated the ratio of one-leaved to three-leaved flowering trillium, and the ratio of vegetative to flowering three-leaved trillium, then, we added one to the ratios and applied a natural log transformation. We used the ratios as responses in separate linear models, with the proportion of flowering trillium symptomatic and total number of flowering trillium in the plot as interacting predictors. Next, we used the absolute abundance of one-leaved and three-leaved vegetative trillium as responses in separate Poisson-distributed generalized linear models (each year tested independently), with the same predictors as described above.

### Geographic distribution of infection

To understand the outcomes of vectored phytoplasma transmission beyond a local population scale, we characterized spatial patterns of phytoplasma infection symptoms of *T. grandiflorum* at a range-wide scale and at range margins. First, we collected data from user observations on iNaturalist, then we annotated all visually symptomatic *T. grandiflorum* observations with the observation field “Phytoplasma Infection?=Yes”. We considered petal virescence and phyllody as the main indicators of phytoplasma infection (Figure 1). While it is possible that these symptoms can arise from other factors, like hormonal imbalances in plants (Mor and Zieslin 1992), or that a plant is infected but asymptomatic (Donkersley et al. 2020), they remain highly characteristic of phytoplasmas and provide a proxy for infection across an otherwise intractable number of plants and populations. On October 7th, 2025, we downloaded all research-grade iNaturalist observations of *T. grandiflorum* from the Global Biodiversity Information Facility (iNaturalist contributors, iNaturalist 2025) and merged them with their associated metadata and observation fields from iNaturalist, downloaded on the same day (data included in supplementary material). After cleaning the dataset with *CoordinateCleaner* (Zizka et al., 2019; Supplemental material), there were 23,218 total observations and 384 symptomatic observations.

Next, we examined patterns of trillium observations across its range, and used Bayesian spatial models to link trillium observation density, as well as other environmental and anthropogenic factors which may affect the ecology of plants and vectors (Santos et al., 2024; Pedroncelli et al., 2026), to the likelihood that a trillium observation displayed evidence of a phytoplasma infection. First, we assigned observations to hexagonal grid cells drawn at a scale of 50 km between opposite sides of the hexagons to quantify trillium observation density across the range of observations. Then, we tested for significant spatial clustering of trillium observations and symptomatic trillium using Moran’s I and the moran.mc function (*spdep*; Pebesma and Bivand, 2023). We visualised grids using the *sf* (Pebesma 2018), *mapview* (Appelhans et al., 2015), and *rnaturalearth* (Massicotte and South, 2017) packages. For each observation, we used the *geodata* package (Hijmans et al., 2025) to extract 30-seconds spatial resolution of worldclim bioclimatic variables, cropland distribution (Zanaga et al., 2021), and human footprint circa 2009 (Venter et al., 2016). Next, we used deterministic Bayesian Integrated Nested Laplace Approximation (INLA) models to determine factors predicting a trillium observation’s symptoms (Rue et al., 2009). INLA models can effectively account for spatial autocorrelation of data and model continuous spatial processes like disease. We used a model selection procedure to exclude predictors that increased model deviance information criterion (DIC) by >2 (*ggregplot*; Albery 2025), but we did not exclude predictors that we had an *a priori* reason to include, such as footprint and observation density. Then, we removed multicollinear variables using vifcor (*usdm*; Naimi et al., 2014), until all predictors had a variance inflation factor below 5. We fit the base model using predictors of cropland, footprint, total trillium number within the observation’s 50 km grid cell, mean diurnal range, temperature seasonality, mean temperature of wettest quarter, mean temperature of driest quarter, mean temperature of warmest quarter, precipitation seasonality, and precipitation of wettest quarter. The binary response variable was the presence or absence of phytoplasma symptoms for each observation. Then, we fit the same model incorporating spatial autocorrelation using a stochastic partial differentiation equation (SPDE) random effect (Lindgren et al., 2011); we set priors following Albery et al. (2020). We constructed the spatial mesh used for prediction across white trillium range by creating a convex hull around the iNaturalist observations, with an outer boundary buffer layer that contained no observations to reduce edge effects (parameters: offset = c(0.5, 1), max.edge = c(1, 5), cutoff = 0.5). When the 2.5% and 97.5% posterior estimates of predictors did not overlap zero, we considered this strong evidence that the predictor affected infection. We used the ggField function (*ggregplot*; Albery 2025) to plot the INLA model spatial field, which represents the marginal posterior mean effects of the spatial random effects on infection status.

## Results

### Identification of potential insect vectors and phytoplasma strain

We amplified a phytoplasma sequence 1,103 base pairs long from symptomatic *T. grandiflorum* tissue at KSR. The strain of phytoplasma most closely matched and had 95.3% similarity to the phytoplasma previously identified in white trillium by Arocha-Rosete et al. (2016), which was related to *Candidatus* Phytoplasma pruni, group 16SrIII-F. In total, we captured 906 insects in the order Hemiptera from the sticky traps, including 14 morphotypes of leafhoppers (Hemiptera: Cicadellidae). Of the 110 insects tested for phytoplasma presence, we found 4 individuals across 2 leafhopper morphotypes that harboured phytoplasma infection at KSR, making them candidates for vectors (Figure 1GH; Figure S4; Table S1). The phytoplasma-positive insect specimens had 96.6-99.1% sequence similarity to leafhoppers in the *Empoasca* genus, which aligns with the phenotype of the morphotypes (Figure 1GH). Whether these species can successfully transmit phytoplasma to trillium hosts has yet to be confirmed experimentally. Further details and raw sequences are provided in the supplement (Table S1; Supplemental file) and on the Sequence Read Archive (Boyle et al., 2026 *b* [BioProject PRJNA1321329]).

### Effects of phytoplasma infection on Trillium grandiflorum growth and reproduction

Trilliums with visible symptoms of phytoplasma infection were smaller and showed reduced reproduction through altered morphology and loss of functional reproductive structures (Figure 1E). Symptomatic trilliums were on average 15.4% shorter than asymptomatic trilliums (*X*^2^ = 8.77, *p* = 0.003; Figure S4, Table S2), a difference of 4.26 cm. While the length of leaves was not significantly different (*X*^2^ = 0.491, *p* = 0.484; Table S2), sepals were on average 28% longer in symptomatic individuals (*X*^2^ = 25.6, *p* < 0.001; Figure S5, Table S2). Symptomatic individuals produced on average 5.85 ± 1.18 seeds while asymptomatic trillium produced an estimated 18.5 ± 0.79 seeds (*X*^2^ = 34.7, *p* < 0.001 ; Figure 2A, Table S3). While this difference is significant, we consider this result exploratory, because seed count in asymptomatic plants was modelled as a function of size, rather than directly measured and counted. 63% of symptomatic trillium produced no seeds.

High frequency of symptomatic individuals reduced the ratio of one-leaved trillium stage classes compared to flowering trillium (*F*_1,138_ = 4.54*, p* = 0.035; Table S4, Figure 2B), perhaps due to reduced seed production caused by infection. We found no significant effect of symptom frequency on the ratio of vegetative to flowering three-leaved trillium in 2023 or 2024 (Table S4). Symptom frequency had a non-significant negative relationship with the absolute number of one-leaved trillium in 2024 and vegetative three-leaved trillium in 2023, however it did significantly reduce the number of vegetative three-leaved trillium in 2024 (*F*_1,138_ = 5.28*, p* = 0.022; Table S5). As expected, flowering trillium significantly increased the absolute abundance of vegetative three-leaved trillium in 2023 (*F*_1,62_ = 228*, p* < 0.001) and 2024 (*F*_1,138_ = 60.3*, p* < 0.001). The absolute number of one-leaved trillium was only affected by the interaction between symptom frequency and number of flowering trillium; flowering trillium counteracted the negative effect of symptom frequency on one-leaved trillium abundance (*F*_1,138_ = 23.6*, p* < 0.001).

### Within-population infection dynamics

Phytoplasma infection symptoms within populations were more likely at low densities of trillium, indicating negative density-dependent dynamics (Figure 3AB). Our survey plots included a wide range of densities (2-74 flowering trillium per 1 m^2^) and symptom prevalence of flowering trillium ranged from 0% to 21% among plots (Figure 3A). We only used flowering trillium to assess putative infections because key symptoms are identifiable by flower inspection. The zero-inflation component of the model indicated that the presence of symptoms was significantly more likely when flowering trillium were at low density (*z*-value = 2.56, *p* = 0.010; Figure 3B, Table S6) and more likely in 2023 compared to 2018 (*z*-value = -2.07, *p* = 0.038). The count component of the model found no significant effect of year (*z*-value = 0.377, *p* = 0.706; Figure 3C, Table S6) or flowering trillium density (*z*-value = 1.81, *p* = 0.069) on the number of symptomatic trilliums in a plot. Thus, both the probability of symptom presence and the proportion of symptomatic individuals was higher at lower densities. When we modelled data from only 2023 and included the prevalence of Hemiptera juveniles on flowering trillium as a predictor in the model, we found that an increased prevalence of Hemiptera juveniles was marginally associated with an increase in the number of symptomatic trillium (*z*-value = 1.86, *p* = 0.063; Table S7). Consistent with our analysis of both years, we found that symptomatic trilliums were more likely to be present in plots with low densities of reproductive stage trillium (*z*-value = 2.22, *p* = 0.026; Table S7). The zero-inflated model characterising symptoms across both sampling years had a better fit with the inclusion of categorical location as a random effect (log likelihood of -254.8 compared to -272.1), but for models using only 2023 data we excluded random effects of categorical location as this improved fit (log likelihood -82.6 compared to - 83.3).

**Figure 3.**
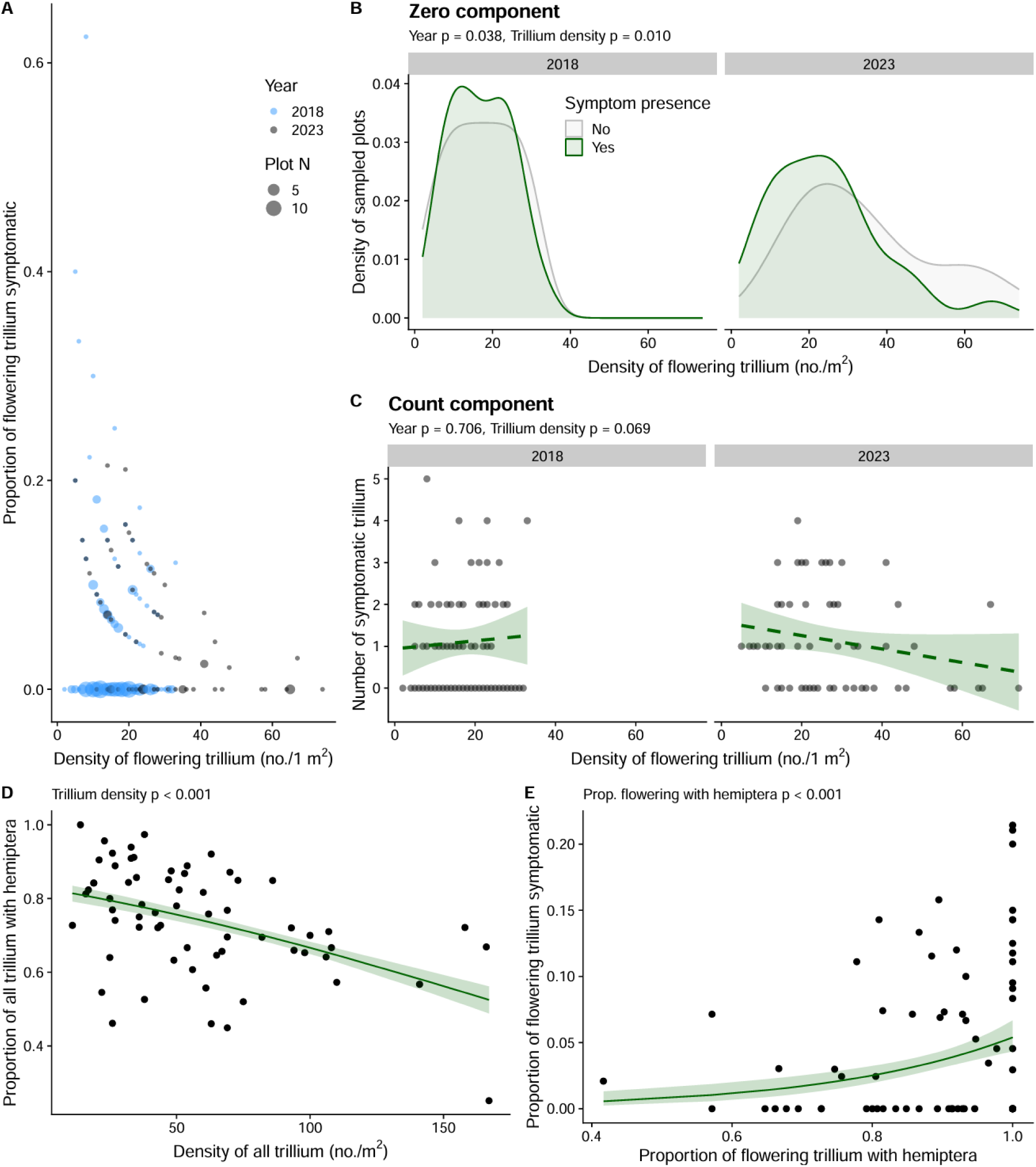
Density-dependent phytoplasma symptom patterns within temperate forest populations at Koffler Scientific Reserve (Ontario, Canada). Panel A is for visualization purposes only and shows a negative relationship between the density and symptoms of flowering trillium, with color denoting year and point size showing the number of field plots with those values. We tested the relationship in (A) using a zero-inflated model (BC). Panel B uses a kernel density estimate to show the density of sampled plots along different trillium densities, coloured by whether there were one or more symptomatic trilliums in the plot; significance values are from the binomial component of a zero-inflated model. Panel C shows the number of symptomatic trilliums across trillium density and year; significance values are from the count component of a zero-inflated model, dashed lines show non-significant trend lines. Panel D shows the relationship between trillium density and the proportion of trillium found with phytophagous Hemiptera juveniles. Panel E shows the proportion of flowering trillium with Hemiptera and the proportion of flowering trillium with symptoms. The scatterplots in D and E show data points and fits from binomial generalised linear models. Alt text: A multi-panel figure showing results from field observations. Collectively the panels show that the likelihood of symptoms is higher in less dense trillium plots, and that Hemiptera occupy more of the trillium plants in less dense trillium plots.

The density of trillium in patches influenced the prevalence of sap-sucking insect juveniles on trillium hosts. As total trillium density increased in a plot, the proportion of total trilliums with Hemiptera juveniles decreased (z-value = -10.786, *p* < 0.001, Figure 3D, Table S8). As the proportion of flowering trillium with Hemiptera juveniles increased in a plot, the proportion of symptomatic trillium also increased (z-value = 4.65, *p* < 0.001, Figure 3E, Table S8). Hemiptera juveniles were highly prevalent and occurred on 42% to 100% of flowering trillium (Figure 3E).

### Geographic distribution of infection

Trillium observations and phytoplasma symptoms were spatially structured and clustered across the landscape (Monte-Carlo simulation of Moran’s I *p* = 0.001 for each, Figure 4AB). Symptomatic trillium observations were more clustered than trillium observations at large (Moran’s I of 0.376 compared to 0.309). Despite clustering, we found observations with phytoplasma symptoms at almost all putative range edges of trillium, even when there were relatively few observations in total (Figure 4AB). The northern range edge showed the least evidence of infection (Figure 4B), with the model predicting low likelihood of symptoms north of 46°N (Figure 4C). The INLA model spatial field, representing the marginal posterior mean effects of the spatial random effects on symptoms, indicated a high likelihood of symptoms across the range of *T. grandiflorum*, especially in the central, southern, and western parts of the range (Figure 4C; standard deviation shown in Figure S6). The spatial model indicated that higher mean temperatures of the warmest quarter were marginally negatively correlated with the likelihood of an observation’s symptoms (mean effect -0.277, credible interval of -0.568 to 0.003) and cropland cover was positively associated with an increased likelihood of an observation’s infection symptoms (mean effect 1.04, credible interval of 0.323 to 1.75) (Figure 4D; Table S9). All other predictors in the spatial model were not associated with likelihood of infection symptoms, including the number of trillium observations nearby and human footprint. All predictors had credible intervals non-overlapping with 0 in the base INLA model without spatial structure, except observation density, precipitation seasonality, and mean temperature of the wettest and driest quarters (Figure 4D; Table S20). INLA model fit was substantially improved by incorporating spatial random effects (ΔDIC = 286).

**Figure 4.**
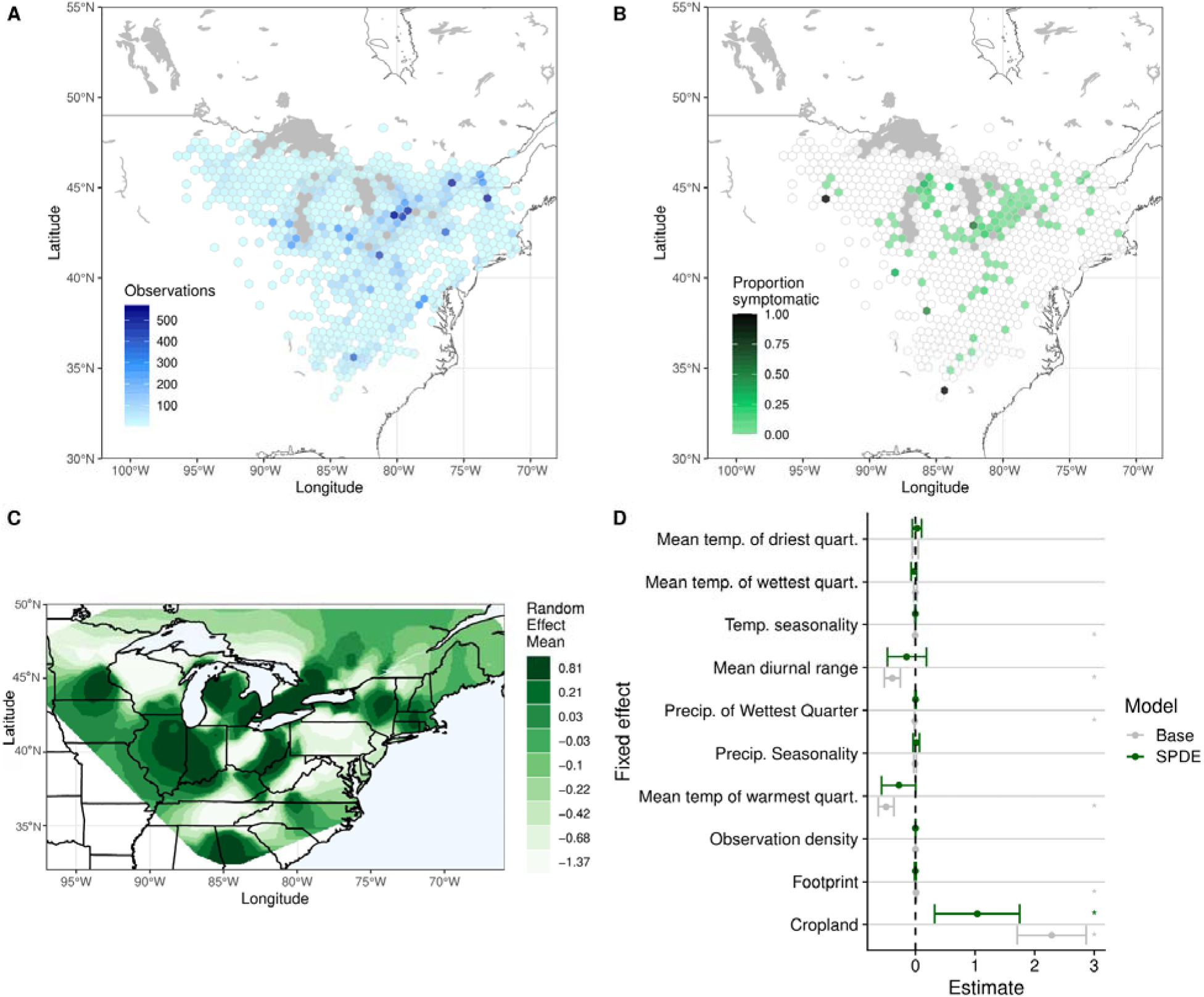
Patterns and predictors of phytoplasma infection symptoms across the range of *Trillium grandiflorum*. The dataset had 23,218 observations (A) with 384 observations of phytoplasma-symptomatic plants (B). The 50 km^2^ hexagonal grid in (A) and (B) represents areas with at least one observation of *T. grandiflorum*; shading in (B) represents the proportion of observations in that hexagon that show symptoms, and white hexagons had no symptomatic observations. The INLA spatial field (C) represents the effect that spatial position had on symptoms accounting for all other covariates in the model. Spatial effects on symptoms were generally positive, particularly in the southern and western parts of the range (C). Cropland cover was significantly associated with increased phytoplasma symptoms respectively in the model that accounted for spatial autocorrelation (SPDE) (D). Asterisks in (D) indicate that the effect and its confidence intervals do not overlap zero in that model; the model is indicated by asterisk colour, with light grey representing the base model and dark green representing the spatial model. Abbreviations are as follows: min. = minimum, temp. = temperature, precip. = precipitation, quart. = quarter. Alt text: A 4-panel figure. The first two panels show the data collected from iNaturalist observations superimposed on a map and demonstrate that symptomatic trillium are found across the range of observations. The remaining panels show results from the INLA spatial model, one being a map with the effect of space on likelihood of infection symptoms and one showing the estimate size of different predictors and whether it overlaps 0.

## Discussion

We studied the impacts of a sterilizing, vector-transmitted pathogen on white trillium populations and patterns of infection across trillium’s range. Within populations, we found that phytoplasma symptom presence was more likely in low host density plots, where sap-sucking insects occupied a higher proportion of trillium. The absolute number of symptomatic trilliums was similar across all plots that had symptoms present, such that low density plots had a greater proportion of symptomatic individuals. Phytoplasma symptoms were widely distributed across trillium’s range and were associated with proximity to cropland. Below, we discuss the ecological processes at work, and their potential consequences for white trillium and other plant populations.

### Patterns and processes of phytoplasma within and across white trillium populations

One possible mechanism driving negative density-dependent patterns within populations is the encounter-dilution effect (Mooring and Hart, 1992; Krebs et al., 2014), wherein vectors may satiate or saturate or are otherwise unable to occupy all hosts at high host densities (Gilbert 2002). Indeed, our results support vector limitation because Hemipteran juveniles occupied a lower proportion of trillium at higher host densities (Figure 3D), and juvenile presence is likely a good direct and indirect proxy for sap-sucking insect visitation and contact, which includes vectors. Given that we observed host plant saturation with sap-sucking juveniles, it is likely that our results underestimate the strength of the encounter-dilution effect between susceptible phytoplasma vectors, phytoplasma, and trillium. As suggested by theory and other recent empirical work (Antonovics et al., 1995; 2023; Bruns et al., 2019 *b*), our results provide support for the idea that the encounter-dilution effect, a density-dependent process, affects the outcome of vectored disease in a natural plant population. It is also possible that infected trillium could transition to dormancy more than non-infected individuals, leading to sparser patches over time where transmission is high, and further compounding the encounter-dilution effect. Higher mortality or regression did not lead to changes in stage class structure, however, which we discuss more below. If phytoplasmas are transmitted in a negative density-dependent manner, they may be more difficult to eradicate from host populations, because efforts to reduce their abundance by removing infected hosts can lead to the very conditions that promote the fastest phytoplasma population growth, ultimately increasing their resilience (Churcher et al., 2006).

The iNaturalist data and the spatial model suggest that this vectored pathogen is likely not constrained to specific regions of the host range, providing evidence that disease can be maintained at both the core and edges (Bruns et al., 2019 *a*). There may be several contributing factors creating this pattern. First, if vectored pathogens are more able to colonize small and low-density host populations compared to other types of pathogens, that may lead to reservoirs of infection that bridge populations. Secondly, while leafhoppers often disperse a few dozen meters a day (Zhou et al., 2003), they can occasionally disperse long distances using wind-mediated migration (Santos et al., 2024), meaning that phytoplasma may be less dispersal-limited compared to aerially or contact transmitted pathogens. Finally, vectored pathogens also benefit from the cognition of their vector, as many insects are experts at seeking out their host plants in the environment (Webster and Cardé, 2017; Jones et al., 2022). At the landscape scale, vectors may find plant hosts based on the frequency of the host relative to other plant species (Antonovics et al., 1995), aligning with processes occurring in frequency-dependent transmission. We did not design our study to explicitly test whether phytoplasma infection at the range scale correlates with trillium density because iNaturalist observation density does not necessarily correlate with trillium population size or density. Regardless, our work suggests that phytoplasma transmission maintains infection at most range edges and motivates more experimental work to understand range-level drivers of vectored pathogens.

Environmental factors were largely non-predictive of characteristic phytoplasma infection symptoms at the range-wide scale. We found that higher mean temperatures of the warmest quarter were marginally correlated with reduced chance of infection, which was unexpected given the hypothesized relationship between heat, vector range, and plant infection susceptibility (Hector et al., 2022; Santos et al., 2024). High heat could correlate with some other aspect of phytoplasma ecology, perhaps through vector identity, activity levels or phenology, or cool areas may coincide with historical regions of disease introduction. Additionally, we found that cropland cover was significantly associated with increased likelihood of infection in *T. grandiflorum*, possibly because phytoplasmas are a major disease in agricultural crops (Rao et al., 2018; Kumari et al., 2019).

### Caveats of field and spatial data

Combining citizen science data and hypothesis-driven field work is an especially powerful way to understand disease dynamics and patterns in plants (Meentemeyer et al., 2015). Using a visual proxy of phytoplasma infection and iNaturalist data allowed us to survey trillium across an otherwise intractable number of plants and locations, however our data has important caveats. While the phenotypes of petal virescence and phyllody are characteristic of phytoplasma infection (Lee et al. 2000), asymptomatic infections are possible and go undetected using visual inspection methods (Donkersley et al. 2020), potentially underestimating the scope and scale of infection within our population and across the range. Asymptomatic infections can only be detected using microscopy, serological tools, or molecular detection tools, which becomes infeasible due to cost and time to implement in ecological surveys on thousands of plants (Donkersley et al. 2020; Bertaccini 2026; Pedroncelli et al., 2026). For example, in our field surveys of infection symptom frequency alone, we inspected 7,141 plants cumulatively across years. In other species, visual symptoms of phyllody and virescence were significantly and reliably predictive of phytoplasma presence, as determined by PCR (Padovan and Gibb 2001; De La Rue et al. 2003). Given that we did not measure asymptomatic infections, it is unknown whether they are common in trillium and if their ecological patterns and drivers differ from symptomatic infections. For example, if prevalences of asymptomatic and symptomatic infections are not tightly correlated, it is possible that by sampling asymptomatic infections we may have seen stronger or weaker negative density-dependence, relationships between stage classes and infection frequency, or relationships between infection probability across the range and environmental predictors. In contrast, if the prevalence of asymptomatic and symptomatic infections are correlated, then results are unlikely to be qualitatively different. Given the interesting patterns we observed at local and range-wide scales, our study motivates further investigation using molecular detection.

The trillium observations showed spatial clustering likely due to the real structure of trillium populations as well as biases like human population density, activity, and location accessibility. Human footprint and the number of trillium observations were weakly positively correlated, however neither were significant predictors in the spatial INLA model, suggesting that the increased sampling effort alone did not change the likelihood of an observation’s infection status. If symptomatic trilliums were more likely to be documented than non-symptomatic trillium, this may have increased the likelihood of phytoplasma detection in populations. Additionally, the full range of abiotic factors experienced by trillium were likely not captured or evenly sampled by our iNaturalist data (Figure S7), due to human bias in location; when some values of the predictor space are more sparsely sampled, this may make it harder to detect relationships between predictors and response, or between predictors. To help reduce these biases, our model selection procedure removed predictors with significant collinearity, making it less likely that our significant geographic predictors are being driven by other predictors such as human footprint and nearby observation number.

Our spatial analysis modelled the chance that an observation was symptomatic, and it provided an estimate of how continuous space affected that probability. While we present the proportion of trillium observations with symptoms over space for visualization purposes (Figure 4B), the proportions are not meant to reflect actual phytoplasma infection frequencies within those populations, nor have we attempted to model infection frequency directly with our model. So, while there are strong limitations to what we can infer with our spatial data, the disease phenotype remains an indicator that phytoplasma may be present across the majority of trillium range. To further test the hypothesis that phytoplasma is range-wide, future research should aim to sample trillium populations across the range systematically for phytoplasma infection, using molecular methods, and accounting for potential biases stemming from iNaturalist-collected data.

### Implications of phytoplasma for plant ecology and evolution

Given the fitness cost accrued with phytoplasma infection (Figure 2A; Lee et al. 2000) and given that leafhoppers transmit the pathogen through feeding, we predict that plant traits involved in herbivore attraction and defense are under selection (Mauricio and Rausher, 1997; Agrawal 2011). However, white trillium does not appear to have many physical defenses against herbivory (Figure 1) and its chemical defense remains unknown. Deer herbivory is known to play a role in structuring trillium populations (Knight 2003; 2007), in contrast, insect herbivory remains largely unstudied (but see Rathcke 1985). It is clear phytoplasma affects fitness components through direct sterilization and conceivably by pollinator avoidance (due to altered floral morphology). Thus, phytoplasma has the potential to change the selective landscape of plants, and more research is needed to understand if there is any subsequent phytoplasma-mediated selection on plant traits in trillium populations, and how this interacts with other important processes such as growth, pollination, and seed dispersal.

### Implications of phytoplasma on wild plant population persistence

Theory predicts high rates of local extinction when pathogens are sterilizing, non-lethal, and in a long-lived species (Anderson and May, 1981; Boots et al., 2002; Gerber et al., 2005; Antonovics 2009). The trillium populations at KSR remain productive after hosting phytoplasma for more than 20 years, though the amount of change in spatial structure, absolute trillium abundance, and infection prevalence over that time remains unknown. At the patch level, only the relative abundance of the one-leaved stage class was correlated with phytoplasma symptoms, but the effect was not large (Figure 2B). We may not see a larger shift in demographic structure because long-lived populations could experience high lag, or because the KSR populations remain large enough that the reduced seed input is not yet materially affecting recruitment rates. Our plots were smaller than the seed dispersal kernel of trillium (Kalisz et al., 1999; Vellend et al., 2003), so it’s likely that plots with high infection receive seed subsidies from surrounding areas. The reduced number of three-leaved vegetative trillium in plots with high infection could suggest that reproductive infected trilliums are dying or regressing to dormant stages at the same rate that younger classes are not being recruited, leading to stable stage classes with an overall reduction in trillium abundance. Entry into dormancy can be instigated by other stressors like deer herbivory (Knight 2003). Another alternative explanation is that trilliums continue to reproduce by rhizomes, masking the demographic consequences of plants producing fewer seeds. However, whether infected trillium continue to produce infected rhizomes is unknown (Lee et al., 2000; Ustun et al., 2023). Seeds produced by symptomatic plants may also carry infection, though more research is needed (Wei and Zhao 2025). Multiple modes of transmission may compound the difficulties small populations may face (Uricchio et al., 2023), while providing a veneer of stability. The full demographic consequences of phytoplasma infection may have yet to materialize. Given that multiple *Trillium* species are hosts of phytoplasma (Arocha-Rosete et al., 2016), populations of conservation concern are the endangered species of trillium that overlap in range with *T. grandiflorum* (Meredith et al., 2022).

