## Supplemental tables and figures for "The ecological dynamics and consequences of phytoplasma infection in the spring ephemeral white trillium"

**Supplemental iNaturalist Methods**

On October 7th, 2025, we downloaded all research-grade iNaturalist observations of *T. grandiflorum* from the Global Biodiversity Information Facility (iNaturalist contributors, iNaturalist 2025) and merged them with their associated metadata and observation fields from iNaturalist, downloaded on the same day (data included in supplementary material). In total, this dataset included 36,497 observations between 2012-2025 with 637 symptomatic individuals. We removed observations with missing latitudes or longitudes, observations with low or missing positional accuracy (uncertainty >10,000 m), and observations likely to be erroneous with both a latitude above 20°N and a longitude below 65°W, resulting in 27,803 observations. To reduce user bias in the dataset, we limited the maximum number of observations per user to 15 by randomly subsampling, reducing the dataset to 24,103 observations. Then, we used *CoordinateCleaner* (Zizka et al., 2019) to remove observations at country centroids, known biological institutions, observations with equal latitudes and longitudes, observations with any zeroes as coordinates, and observations with identical coordinates. Users contributed a mean 1.89 total observations and a median of 1 total observation. The symptomatic observations were observed by 319 users. We reprojected the coordinates of observations using ESRI:102003, an equal area projection that is appropriate for *T. grandiflorum*’s range.

**Supplemental figures**

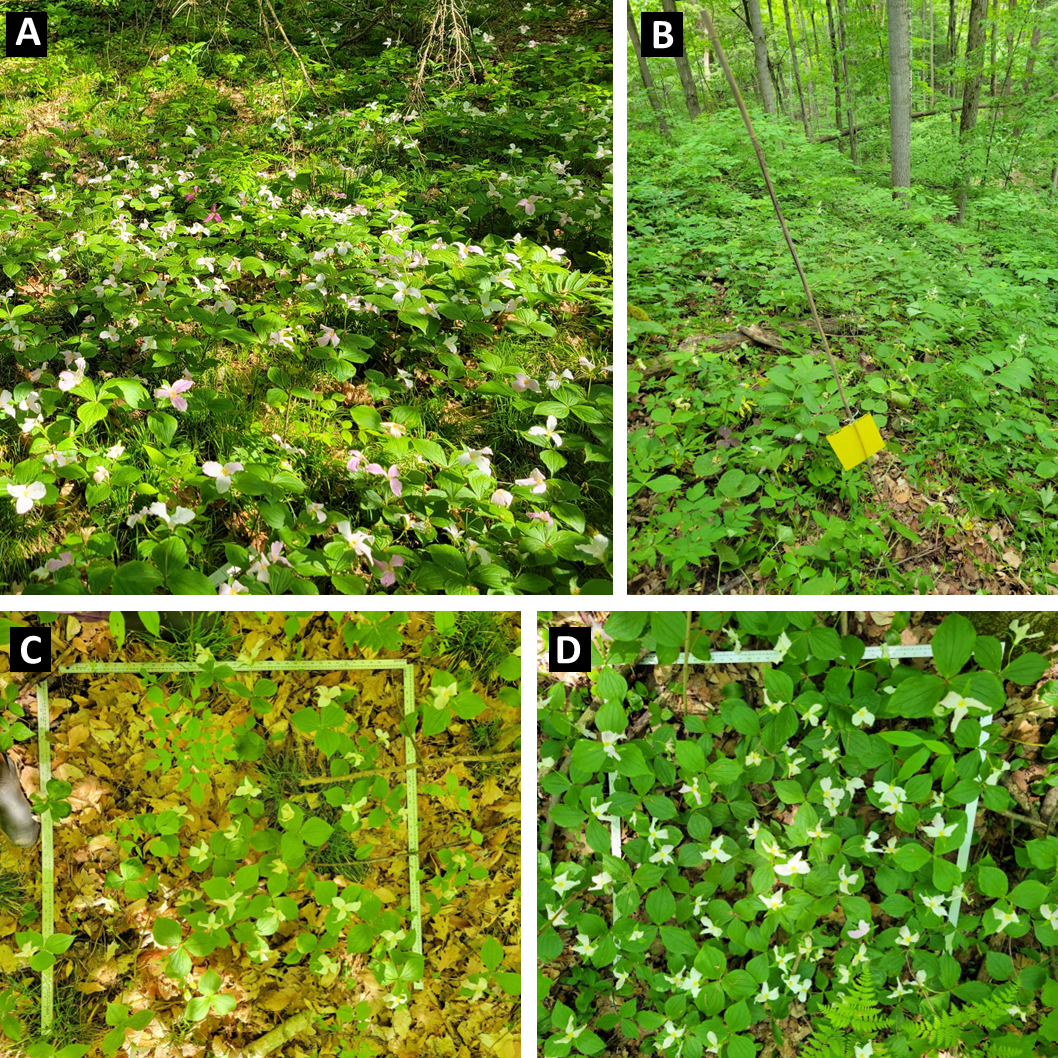

Figure S1. White trillium at the Koffler Scientific Reserve and methods used to sample. White trillium is a common spring ephemeral at the field station (A). We placed sticky traps for trapping and sequencing leafhopper vectors (B). We sampled 1 m^2^ plots of varying trillium density (C, D). Photos taken by authors in 2023.

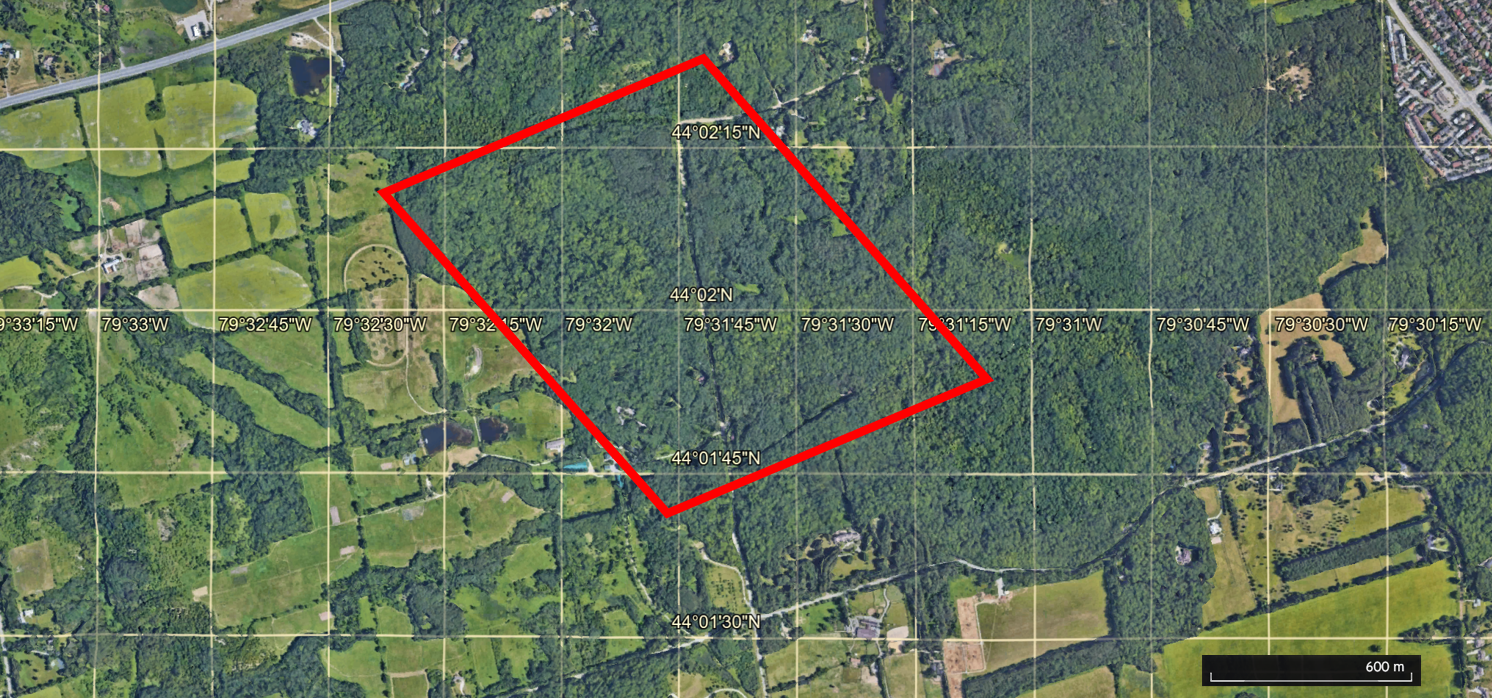

Figure S2. Location of the white trillium population at the Koffler Scientific Reserve is indicated by the red rectangle. Subsamples of this population were performed in 2018 and 2023. Marked grid lines display latitude and longitude, and a scale bar is in the lower right-hand corner.

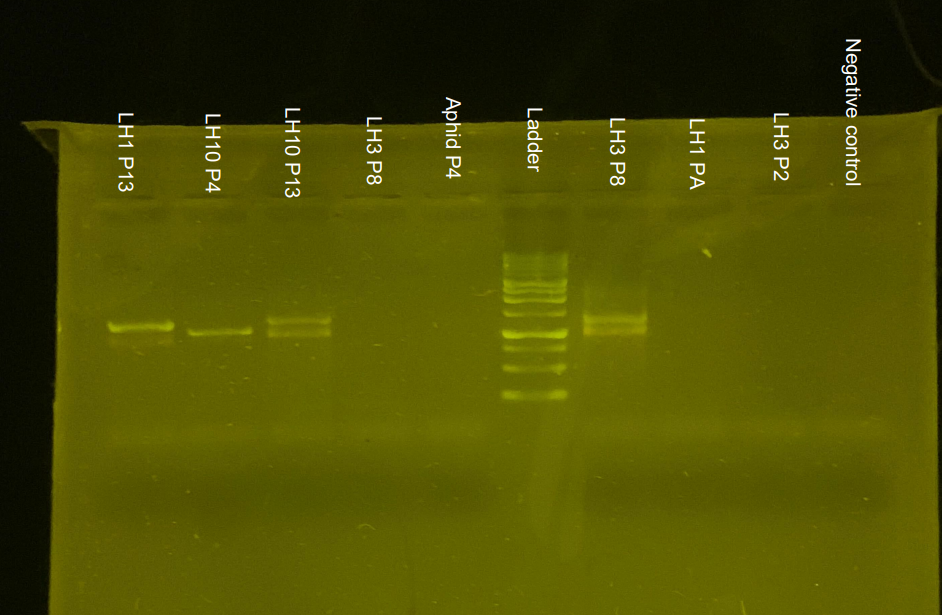

Figure S3. Subset of tested insect samples, containing all positive samples. LH = Leaf hopper. The number after LH is the morphotype; we determined morphotypes 1 and 3 were the same after data collection. P# denotes the plot the insect was collected from (all insects also have unique individual numbers, not included here for brevity).

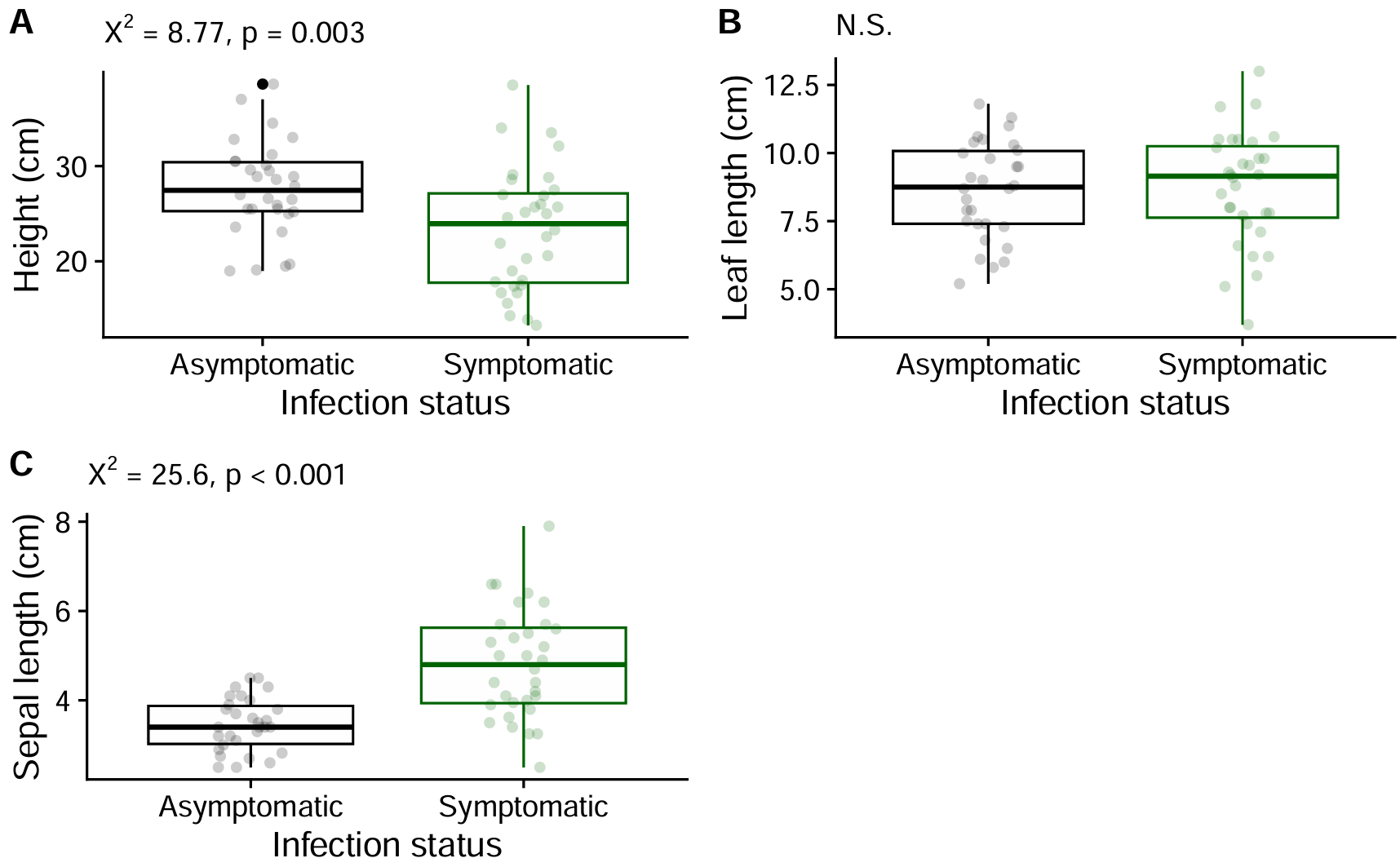

Figure S4. Phenotypic traits of *T. grandiflorum* are altered when symptomatic with phytoplasma. We used petal virescence and phyllody as symptom proxies of phytoplasma infection. Linear mixed models found that height was significantly reduced and sepal length was significantly increased in symptomatic individuals. Leaf length was not significantly different. Data from 2023 surveys (asymptomatic n = 30, symptomatic n = 32).

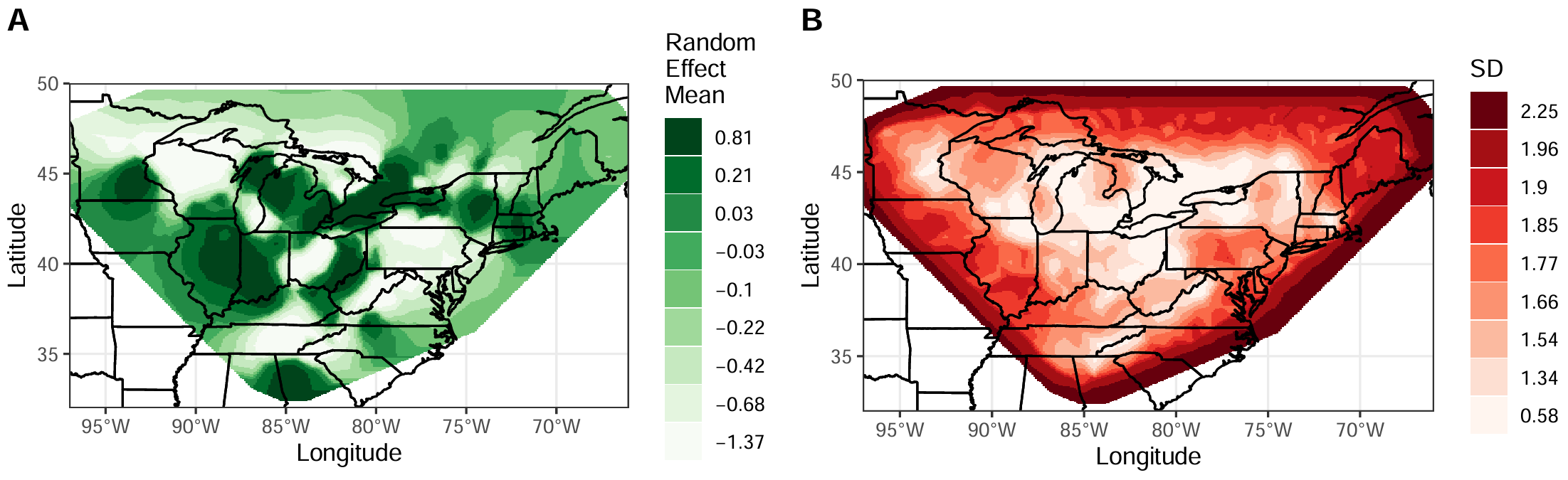

Figure S5. Spatial field mean (A) and standard deviation (B), estimated from the INLA model. Spatial fields represent marginal posterior effects of the random effects of space. Values are shown across the mesh.

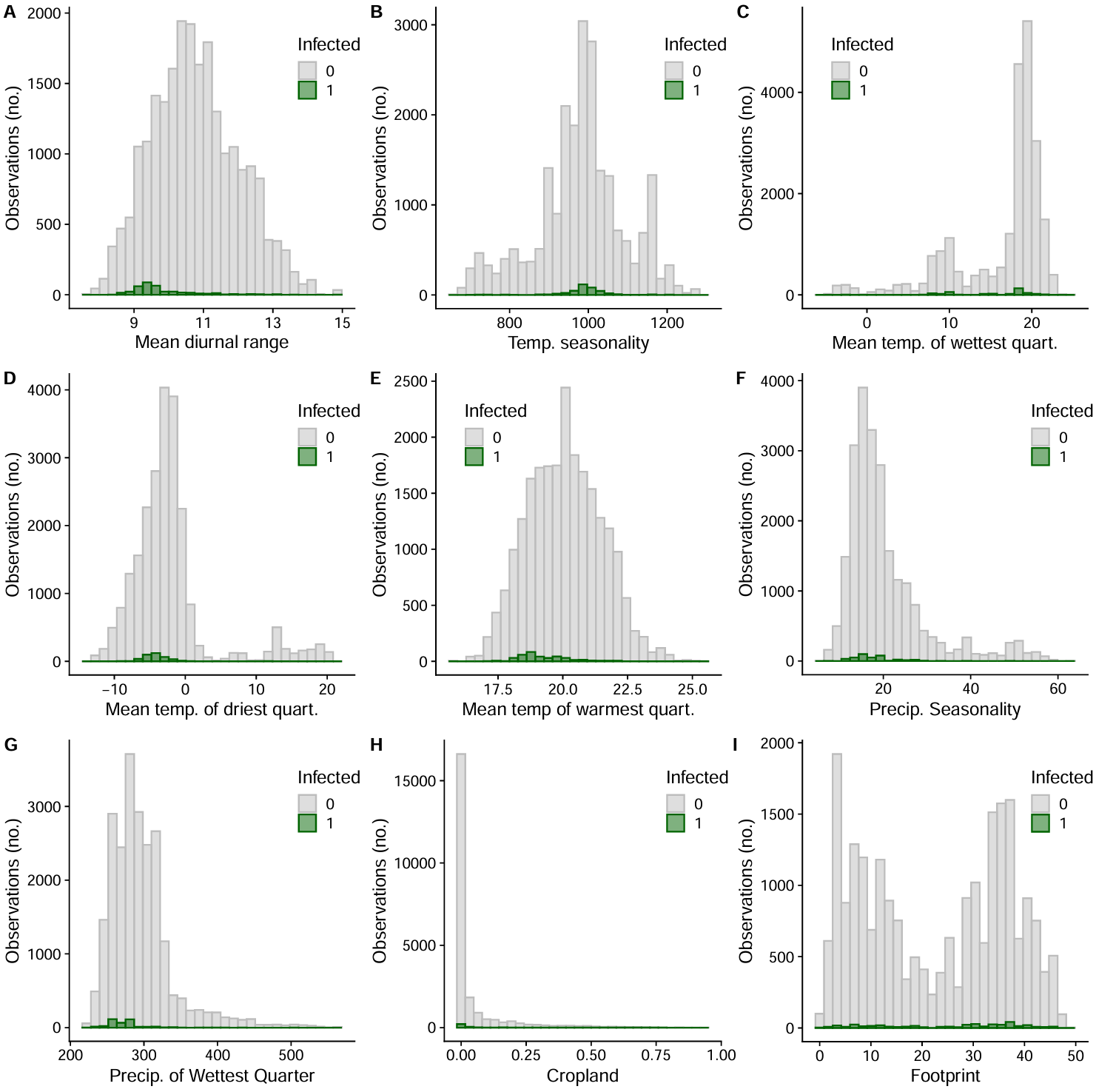

Figure S6. Histograms of the number of symptomatic and asymptomatic observations across all model predictors. Not all values of the predictors are sampled to the same depth--an inherent feature of community science data. If the likelihood and frequency of infection symptoms in populations doesn’t depend on the predictor, then we would expect symptomatic and asymptomatic observations to share the same distribution.

**Supplemental Tables**

Table S1. Number of Hemiptera morphotypes caught and tested, and their phytoplasma positivity rate. Phytoplasma presence was determined with a nested polymerase chain reaction using two universal phytoplasma primer pairs R16mF2/R1 and R16F2n/R2 (Gundersen & Lee 1996), and we visualised the presence of phytoplasma using gel electrophoresis. We determined leafhopper morphotypes 1 and 3 were likely the same species by morphology, but differing in sample freshness/time since mortality.

| **Morphotype** | **Number caught** | **Number tested** | **Number positive** | **Percent positive %** |
| --- | --- | --- | --- | --- |
| Leafhopper 1+3 | 279+223 | 48+20 | 1+1 | 2.94 |
| Leafhopper 2 | 154 | 7 | 0 | 0 |
| Leafhopper 4 | 16 | 1 | 0 | 0 |
| Leafhopper 5 | 6 | 2 | 0 | 0 |
| Leafhopper 6 | 2 | 1 | 0 | 0 |
| Leafhopper 7 | 6 | 0 | NA | NA |
| Leafhopper 8 | 18 | 4 | 0 | 0 |
| Leafhopper 9 | 16 | 2 | 0 | 0 |
| Leafhopper 10 | 58 | 5 | 2 | 40 |
| Leafhopper 11 | 9 | 0 | NA | NA |
| Leafhopper 12 | 4 | 1 | 0 | 0 |
| Leafhopper 13 | 6 | 0 | NA | NA |
| Leafhopper 14 | 1 | 0 | NA | NA |
| Leafhopper 15 | 6 | 0 | NA | NA |
| Leafhopper nymph 1 | 3 | 1 | 0 | 0 |
| Leafhopper nymph 2 | 3 | 3 | 0 | 0 |
| Leafhopper nymph 3 | 1 | 1 | 0 | 0 |
| Treehopper 1 | 2 | 0 | NA | NA |
| Treehopper 2 | 2 | 0 | NA | NA |
| Psyllid 1 | 11 | 9 | 0 | 0 |
| Psyllid 2 | 1 | 0 | NA | NA |
| Aphid 1 (Black colour) | 49 | 3 | 0 | 0 |
| Aphid 2 (Long legs) | 2 | 0 | NA | NA |
| Aleyrodidae/whiteflies | 28 | 0 | NA | NA |
| **TOTAL** | **906** | **110** | **4** | **3.64** |

Table S2. Phenotypic response of trillium to phytoplasma symptoms. Data from 2023 surveys (asymptomatic n = 30, symptomatic n = 32). Estimates are relative to asymptomatic plants.

| **Predictor** | **Height (cm)** | | | | **Leaf length (cm)** | | | | **Log of Sepal length (cm)** | | | |
| --- | --- | --- | --- | --- | --- | --- | --- | --- | --- | --- | --- | --- |
|  | *Estimate* | *Std. Error* | *Wald Chisq* | *p* | *Estimate* | *Std. Error* | *Wald Chisq* | *p* | *Estimate* | *Std. Error* | *Wald Chisq* | *p* |
| Intercept | 27.8 | 1.12 | 611 | **< 0.001** | 9.09 | 0.525 | 300 | **< 0.001** | 1.30 | 0.063 | 430 | **< 0.001** |
| Infection status | -4.39 | 1.48 | 8.77 | **0.003** | -0.342 | 0.488 | 0.491 | 0.484 | 0.282 | 0.056 | 25.6 | **< 0.001** |

Table S3. Phytoplasma symptoms effects on white trillium fitness, accounting for trillium size and location. Infection status estimates are relative to asymptomatic plants, and modeled seed count status is relative to predicted seed counts. Boxes are greyed if not relevant to the model under ‘model goal’.

| *Response* | **Seed count** | | | | | | | |
| --- | --- | --- | --- | --- | --- | --- | --- | --- |
| *Model goal* | Do symptomatic trillium make fewer seeds than predicted based on their size? | | | | Do symptomatic trillium make fewer seeds than a nearby asymptomatic trillium? | | | |
| *Predictor* | *Estimate* | *Std. Error* | *Wald Chisq* | *p* | *Estimate* | *Std. Error* | *Wald Chisq* | *p* |
| Intercept | 2.77 | 0.119 | 541 | **< 0.001** | 2.91 | 0.116 | 635 | **< 0.001** |
| Modeled seed count status | -1.21 | 0.216 | 31.3 | **< 0.001** |  |  |  |  |
| Infection status |  |  |  |  | -1.24 | 0.210 | 34.7 | **< 0.001** |

Table S4. How symptom frequency and trillium density affect the ratio of non-reproductive to reproductive stage classes. Estimates are from linear models, while the F values and p values are from type III Anovas applied to the models using the car package.

|  | **log(No. 3-leaved vegetative trillium:Flowering no. in 2023)** | | | | **log1p(No. 3-leaved vegetative trillium:Flowering no. in 2024)** | | | | **log1p(No. 1-leaved trillium:Flowering no. in 2024)** | | | |
| --- | --- | --- | --- | --- | --- | --- | --- | --- | --- | --- | --- | --- |
| **Predictor** | *Estimate* | *Std. Error* | *F* | *p* | *Estimate* | *Std. Error* | *F* | *p* | *Estimate* | *Std. Error* | *F* | *p* |
| Intercept | -0.172 | 0.270 | 0.406 | 0.526 | 1.13 | 0.103 | 121 | **<0.001** | 0.649 | 0.085 | 58.1 | **<0.001** |
| Symptom frequency in flowering trillium | 3.49 | 3.27 | 1.14 | 0.291 | -0.359 | 0.201 | 3.20 | 0.076 | -0.354 | 0.166 | 4.54 | **0.035** |
| Total no. flowering trillium | 0.001 | 0.007 | 0.016 | 0.901 | -0.068 | 0.032 | 4.44 | **0.037** | -0.072 | 0.027 | 7.37 | **0.007** |
| Symptoms : Flowering trillium no. | -0.237 | 0.158 | 2.25 | 0.138 | 0.009 | 0.278 | 0.001 | 0.975 | 0.333 | 0.230 | 2.10 | 0.150 |

Table S5. How symptom frequency and trillium density affect plants in non-reproductive stage classes. Estimates are from the generalized linear models with a Poisson distribution, while the Likelihood Ratio Chi-Square values and p values are from type III Anovas applied to the glm models using the car package.

|  | **Abundance of 3-leaved vegetative trillium in 2023** | | | | **Abundance of 3-leaved vegetative trillium in 2024** | | | | **Abundance of 1-leaved trillium in 2024** | | | |
| --- | --- | --- | --- | --- | --- | --- | --- | --- | --- | --- | --- | --- |
| **Predictor** | *Estimate* | *Std. Error* | *LR Chisq* | *p* | *Estimate* | *Std. Error* | *LR Chisq* | *p* | *Estimate* | *Std. Error* | *LR Chisq* | *p* |
| Intercept | 2.69 | 0.068 | - | **-** | 1.23 | 0.064 | - | **-** | 0.388 | 0.115 | - | **-** |
| Symptom frequency in flowering trillium | -0.693 | 0.954 | 0.533 | 0.465 | -0.387 | 0.176 | 5.28 | **0.022** | -0.340 | 0.268 | 1.72 | 0.189 |
| Total no. flowering trillium | 0.023 | 0.001 | 228 | **<0.001** | 0.128 | 0.015 | 60.3 | **<0.001** | 0.043 | 0.033 | 1.55 | 0.213 |
| Symptoms : Flowering trillium no. | -0.052 | 0.042 | 1.47 | 0.226 | 0.385 | 0.193 | 3.77 | 0.052 | 1.26 | 0.245 | 23.6 | **<0.001** |

Table S6. Zero-inflated model for the population dynamics of the multiyear dataset (2018 and 2023). This model included random effects of sub-population at the Koffler Scientific Reserve. Plots were 1 m^2^.

|  | **Symptomatic trillium presence (zero model)** | | | | **Symptomatic trillium no. (count model)** | | | |
| --- | --- | --- | --- | --- | --- | --- | --- | --- |
| **Predictor** | *Estimate* | *Std. Error* | *z-value* | *p* | *Estimate* | *Std. Error* | *z-value* | *p* |
| Intercept | -0.434 | 0.880 | -0.494 | 0.621 | -0.139 | 0.226 | -0.613 | 0.540 |
| Reproductive trillium no. | 0.064 | 0.025 | 2.56 | **0.010** | 0.016 | 0.009 | 1.81 | 0.069 |
| Year | -3.17 | 1.54 | -2.07 | **0.039** | 0.085 | 0.226 | 0.377 | 0.706 |

Table S7. Zero-inflated model for the population dynamics of the 2023 dataset. This model excluded random effects of sub-population at the Koffler Scientific Reserve. Plots were 1 m^2^.

|  | **Symptomatic trillium presence (zero model)** | | | | **Symptomatic trillium no. (count model)** | | | |
| --- | --- | --- | --- | --- | --- | --- | --- | --- |
| **Predictor** | *Estimate* | *Std. Error* | *z-value* | *p* | *Estimate* | *Std. Error* | *z-value* | *p* |
| Intercept | -10.1 | 8.43 | -1.20 | 0.231 | -2.29 | 1.38 | -1.66 | 0.097 |
| Reproductive trillium no. | 0.099 | 0.044 | 2.22 | **0.026** | 0.014 | 0.012 | 1.11 | 0.267 |
| Proportion of reproductive trillium with Hemiptera nymphs | 6.43 | 7.71 | 0.834 | 0.404 | 2.51 | 1.35 | 1.86 | 0.063 |

Table S8. Binomial generalised linear models testing whether trillium density predicts Hemiptera prevalence (A), and whether Hemiptera prevalence predicts symptom proportions (B).

| **Predictor** | *Estimate* | *Std. Error* | *z-value* | *p* |
| --- | --- | --- | --- | --- |
| A. Proportion of trillium that had Hemiptera on them  ~ number of trillium in a plot | | | | |
| Intercept | 1.57 | 0.080 | 19.77 | <0.001 |
| Number of trillium | -0.00883 | 0.000819 | -10.78 | <0.001 |
| B. Proportion of adult trillium with symptoms of infection  ~ proportion of adult trillium in a plot with Hemiptera | | | | |
| Intercept | -6.82 | 0.780 | -8.74 | <0.001 |
| Proportion with Hemiptera | 3.95 | 0.850 | 4.65 | <0.001 |

Table S9. Posterior summaries of the integrated nested Laplace approximation (INLA) model results with stochastic partial differential equation random effects to account for spatial autocorrelation. The deviance information criterion (DIC) was 3246.76, saturated DIC was 3213.21.

| **Fixed effects** | **Mean** | **Standard deviation** | **0.025 quantile** | **0.5 quantile** | **0.975 quantile** | **Mode** |
| --- | --- | --- | --- | --- | --- | --- |
| Intercept | 2.90 | 4.29 | -5.51 | 2.89 | 11.2 | 2.89 |
| Trillium no. in grid | 0.000 | 0.001 | -0.001 | 0.000 | 0.001 | 0.000 |
| Mean diurnal range | -0.149 | 0.167 | -0.470 | -0.151 | 0.185 | -0.151 |
| Temp. seasonality | -0.002 | 0.003 | -0.007 | -0.002 | 0.004 | -0.002 |
| Mean temp. of wettest quart. | -0.026 | 0.024 | -0.072 | -0.026 | 0.021 | -0.026 |
| Mean temp. of driest quart. | 0.025 | 0.039 | -0.051 | 0.025 | 0.103 | 0.025 |
| Mean temp of warmest quart. | -0.277 | 0.145 | -0.568 | -0.275 | 0.003 | -0.275 |
| Precip. seasonality | 0.013 | 0.027 | -0.039 | 0.013 | 0.067 | 0.013 |
| Precip. of wettest quart. | 0.002 | 0.005 | -0.007 | 0.002 | 0.011 | 0.002 |
| Cropland cover | 1.04 | 0.363 | 0.323 | 1.04 | 1.75 | 1.04 |
| Footprint | -0.001 | 0.007 | -0.014 | -0.001 | 0.012 | -0.001 |
| **Random effects** |  | | | | | |
| Range for spatial effects | 1.78 | 0.447 | 1.09 | 1.72 | 2.84 | 1.60 |
| St. dev. for spatial effects | 1.75 | 0.226 | 1.35 | 1.74 | 2.24 | 1.71 |

Table S10. Posterior summaries of the base integrated nested Laplace approximation (INLA) model results (no stochastic partial differential equation). The deviance information criterion (DIC) was 3532.81, and the saturated DIC was 3499.26.

| **Fixed effects** | **Mean** | **Standard deviation** | **0.025 quantile** | **0.5 quantile** | **0.975 quantile** | **Mode** |
| --- | --- | --- | --- | --- | --- | --- |
| Intercept | 17.5 | 1.73 | 14.2 | 17.5 | 20.9 | 17.5 |
| Trillium no. in grid | 0.001 | 0.000 | 0.000 | 0.001 | 0.002 | 0.001 |
| Mean diurnal range | -0.388 | 0.069 | -0.523 | -0.388 | -0.253 | -0.388 |
| Temp. seasonality | -0.005 | 0.001 | -0.007 | -0.005 | -0.003 | -0.005 |
| Mean temp. of wettest quart. | 0.000 | 0.013 | -0.026 | 0.000 | 0.025 | 0.000 |
| Mean temp. of driest quart. | -0.003 | 0.025 | -0.053 | -0.003 | 0.047 | -0.003 |
| Mean temp of warmest quart. | -0.492 | 0.067 | -0.624 | -0.492 | -0.361 | -0.492 |
| Precip. seasonality | -0.006 | 0.011 | -0.027 | -0.006 | 0.015 | -0.006 |
| Precip. of wettest quart. | -0.012 | 0.002 | -0.016 | -0.012 | -0.007 | -0.012 |
| Cropland cover | 2.29 | 0.295 | 1.71 | 2.29 | 2.87 | 2.29 |
| Footprint | 0.011 | 0.005 | 0.001 | 0.011 | 0.022 | 0.011 |
